# High-resolution melt curve analysis for rapid detection of rifampicin resistance in *mycobacterium tuberculosis*: A single-center study in Iran

**DOI:** 10.1101/754093

**Authors:** Samaneh Arefzadeh, Mohammad Javad Nasiri, Taher Azimi, Zahra Nikpor, Hossein Dabiri, Farahnoosh Doustdar, Hossein Goudarzi, Mohammad Alahyar

**Affiliations:** Department of Microbiology, School of Medicine, Shahid Beheshti University of Medical Sciences, Tehran, Iran; Department of Microbiology, School of Medicine, Tehran University of Medical Sciences, Tehran, Iran; Regional Tuberculosis Reference laboratory, Tehran University of Medical Sciences, Tehran, Iran

**Keywords:** Tuberculosis, high resolution melting analysis, Xpert MTB/RIF, rifampicin resistance, Iran

## Abstract

**Introduction:** Tuberculosis (TB) remains a leading cause of death worldwide, especially in developing countries. Early detection of resistance is extremely important to reduce the risk of death. This study was aimed to compare the diagnostic accuracy of high resolution melting (HRM) analysis in comparison with Xpert MTB/RIF as well as conventional drug susceptibility testing (DST) for the detection of rifampicin (RIF) resistance in *Mycobacterium tuberculosis* in Iran.

**Materials and methods:** A comparative cross-sectional study was carried out from April 2017 to September 2018. A total of 80 culture-positive clinical samples selected during the study period were analyzed for detection of RIF-resistant TB by conventional DST, Xpert MTB/RIF, and sequencing. Sensitivity and specificity of the HRM calculated according to DST as our gold standard test in this study.

**Results:** The overall sensitivity, specificity, positive predictive value (PPV) and negative predictive value (NPV) of HRM assay were found to be 100%, 89.33%, 38.46%, and 100% respectively.

**Conclusions:** The analysis has demonstrated that the diagnostic accuracy of HRM tests is insufficient to replace Xpert MTB/RIF and conventional DST. HRM test may be used in combination with culture due to the advantage of the time to result. Further work to improve molecular tests would benefit from standardized reference standards and the methodology.

## Introduction

Tuberculosis (TB) is one of the most serious public health problems worldwide. Approximately 10.4 million people were infected with TB in 2016, 90% of them were adults [1]. The emergence of multidrug-resistant TB (MDR-TB) has had a significant negative effect on TB control strategies. In 2016, there were 600,000 new cases with resistance to rifampicin (RIF), of which 490,000 had MDR-TB [1]. In Iran as a country with a moderate TB incidence, it was reported that MDR-TB accounted for 12% of previously treated TB cases and 1% of new TB cases [2]. MDR-TB has been associated with worse treatment outcomes than drug-susceptible TB [3–8]. RIF-resistance is frequently associated with concomitant isoniazid-resistance and thus is considered as a proxy for MDR-TB [9]. Thus, for better management of drug-resistant TB, early detection of RIF-resistance and starting sufficient treatment is extremely important to reduce the risk of death [10–13]. Although the conventional drug susceptibility testing (DST) method is the gold standard test to diagnose drug-resistant TB, it is a time-consuming process that takes up to several weeks to show results [14]. The Xpert MTB/RIF is an automated molecular assay endorsed by the WHO for the early diagnosis of both *M*. *tuberculosis* infection and RIF resistance [15]. The assay has been in use in Iran for detection of RIF-resistance. However, implementation of this method in all regional laboratories of Iran is not affordable. High resolution melting (HRM) curve analysis is a simple and robust method for detecting drug-resistant TB [16–21]. The advantages of HRM include rapid turn-around times, a closed-tube method that greatly reduces contamination risk and, unlike other methods, requires no post-PCR handling [22, 23]. The method is based on the analysis of fluorescence curves produced by DNA-binding dye during strand dissociation events in the melting phase following real-time PCR [23–25]. In the present study, we aimed to compare the diagnostic accuracy of HRM in comparison with Xpert MTB/RIF and DST for the detection of RIF resistance in *M*. *tuberculosis* in Iran.

## Materials and methods

### M. tuberculosis clinical isolates

Twenty clinical isolates of *M. tuberculosis* (10 RIF-resistant TB and 10 drug-susceptible isolates) were selected as the reference samples for the initial development of the HRM assay (the specimen used were culture isolates). To validate the HRM assay, 80 clinical isolates (5 RIF-resistant TB and 75 drug-susceptible isolates) which were collected between April 2017 and March 2018 from the regional TB reference laboratory of Tehran, Iran were used for the screening in a blind manner. The reference strain H37Rv, susceptible to RIF, was included in each run as wild-type positive control and nuclease-free water was used as a negative control. The ethics committee of Shahid Beheshti University of Medical Sciences approved the study, and all patients provided written informed consent (IR.SBMU. MSP.REC.1396.107).

### Identification of M. tuberculosis

Clinical specimens were processed by the standard sodium hydroxide method and smears were prepared by the Ziehl-Neelsen staining method [26]. After decontamination, specimens were inoculated to Lowenstein-Jensen (LJ) solid medium. For identification of mycobacteria, the slope cultures were incubated at 37°C and examined for growth once weekly up to 6 weeks. Bacterial isolates identified as *M. tuberculosis* using standard biochemical tests (i.e. production of niacin, nitrate reduction, catalase) and molecular method (*IS6110* based PCR assay) [26, 27]. Only one culture isolated per study subject was considered for further analysis.

### Conventional DST of M. tuberculosis

Conventional DST for RIF was performed for all culture-positive samples with the proportion method on LJ solid medium with a standard critical concentration of 40 μg/ml for RIF as previously described [26]. *M*. *tuberculosis* H37Rv strain (ATCC 27294) was used for quality control testing in DST.

### Xpert MTB/RIF assay

Xpert MTB/RIF procedure was used according to the manufacturer’s instructions for the molecular detection of RIF resistance [15]. Briefly, Xpert sample reagent was added to 1ml of specimens in the ratio 1:2, and then the mixture was transferred to the Xpert test cartridge. Cartridges were inserted into the Xpert machine, and the automatically generated results were read after 90 min.

### DNA extraction

DNA from the clinical isolates of *M*. *tuberculosis* was extracted from bacterial colonies grown on LJ slants as described previously [28].

### HRM assay

The EvaGreen^®^ Master kit (Metabion, Martinsried, Germany) was used for HRM assay containing the following components per reaction mixture: 10 μl of the 5X master mix, 2 μl of the template, 1 μl each primer and PCR grade water adjusted to a final volume of 20 μl. The RIF primer used for this experiment were previously reported [17]. All assays were run on the Rotor-Gene 6000 instrument (Qiagen, USA) using the following cycling parameters: initial denaturation at 95 °C for 10 min, 40 cycles of 94 °C for the 30s and 60 °C for 30s. When a simple PCR amplification was complete, the HRM analysis was performed from 80–94 °C with 0.1 °C increments every 2s. The reference strain H37Rv, susceptible to RIF, was included in each run as wild-type positive control and nuclease-free water was used as a negative control. Rotor-Gene 6000 Series Software 1.7 was used for HRM analysis.

### DNA sequencing

Sequencing was used to confirm resistance in all phenotypically RIF-resistant strains using the primers which have been previously reported [24].

### Statistical analysis

Data were analyzed with MedCalc 14 statistical software. Sensitivity, specificity, positive predictive value (PPV) and negative predictive value (NPV) of the HRM calculated according to DST as our reference standard in this study.

## Results

### DST

Resistance and susceptible isolates were confirmed by both Xpert MTB/RIF and conventional DST. Thus, the concordance between the Xpert MTB/RIF and conventional DST for RIF resistance was 100%.

### Initial development of HRM assay for detection of RIF resistance

For the initial assay development, we tested the assay for mutations within the *rpoB* gene from 10 RIF-resistant *M*. *tuberculosis* isolates. All RIF-resistant *M*. *tuberculosis* isolates were detected by our HRM assay. We also analyzed 10 drug-susceptible isolates to check for false-positive results. All 10 isolates were correctly identified as wild type. The normalized melt curves of PCR products of isolates with different mutations are shown in Figure 1. Visually, samples with mutations (red line) are easily differentiated from the wild type by the distinct differences in the shape of the melt curves.

**Figure1.**
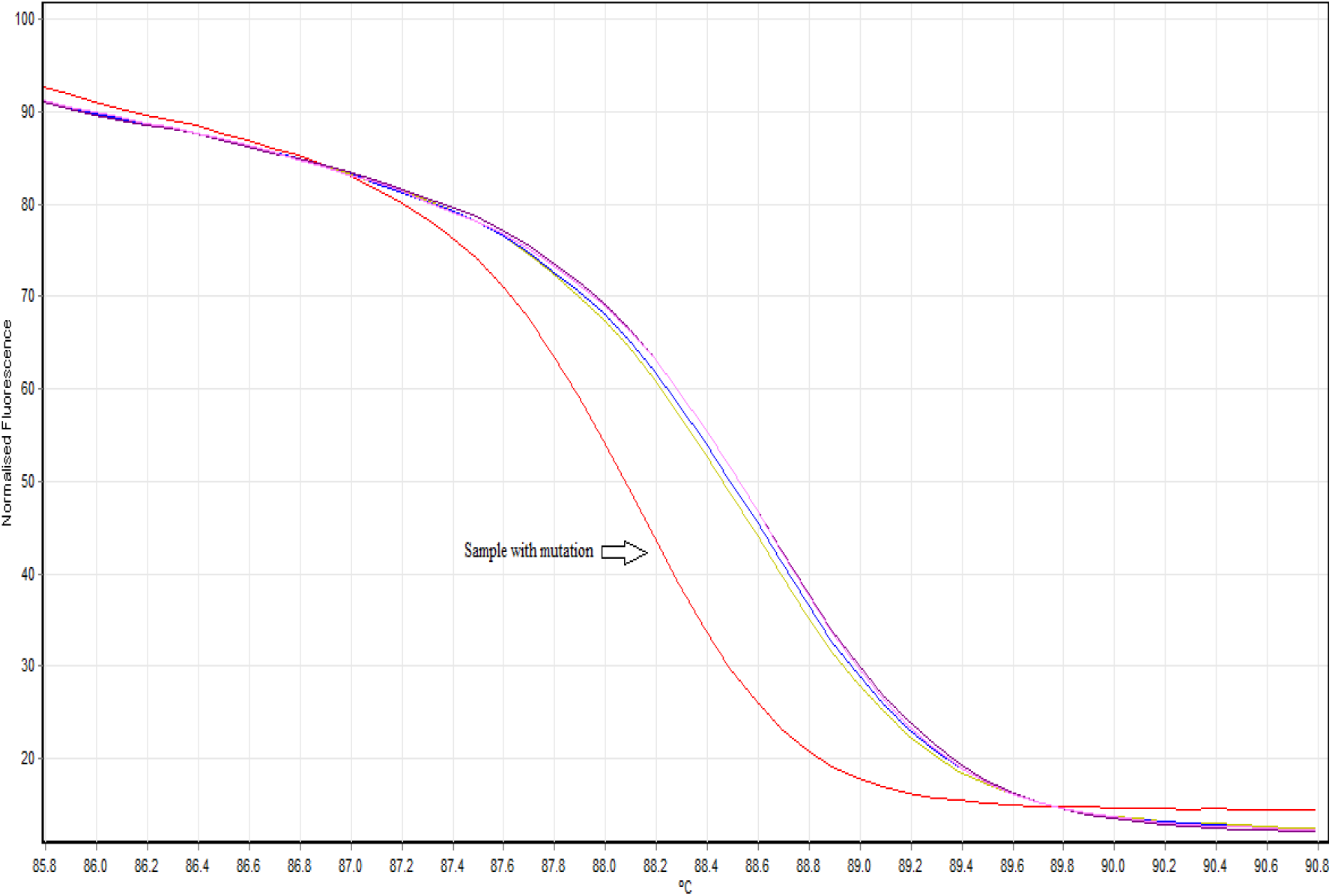
Normalized melting curves of *rpoB*, illustrating the HRM profile separation between wild type strains and strains containing mutations (red).

### Validation of the HRM assay with a blinded series of strains

All 5 RIF-resistant isolates in our blinded samples were detected to have mutations in *rpoB* by HRM (Table1). However, 8 susceptible isolates (confirmed by Xpert MTB/RIF and conventional DST) were falsely clustered as resistance by HRM. These isolates were confirmed for lacking any mutation in the studied regions of *rpoB* by sequencing.

**Table1.** Performance of HRM as compared to conventional drug susceptibility testing.

| HRM | Drug Susceptibility Test (rifampicin)* |  |  | Sensitivity (95% CI) | Specificity (95% CI) |
| --- | --- | --- | --- | --- | --- |
|  | Resistance (5) | Sensitive (75) | Total |  |  |
| Resistance | 5 | 8 | 13 | 100.0 (50.0-100.0) | 89.3 (80.0-95.2) |
| Sensitive | 0 | 67 | 67 |  |  |
| Total | 5 | 75 | 80 |  |  |

345 \* The concordance between the Xpert MTB/RIF and conventional DST for RIF resistance was 100%.

### DNA sequencing

Single-point mutations in a 118 bp region of the *rpoB* gene were observed in all 5 phenotypically resistant isolates.

### Sensitivity and specificity

In order to evaluate the sensitivity and specificity, the HRM curve analysis was compared to the Xpert MTB/RIF and conventional DST results for the detection of RIF-resistant *M. tuberculosis* isolates. The overall sensitivity, specificity, PPV and NPV of HRM assay were found to be 100%, 89.33%, 38.46%, and 100% respectively.

## Discussion

In the current study, we have evaluated the HRM for detection of RIF resistance in *M*. *tuberculosis*. We found that the sensitivity of the HRM was slightly higher than the results found in previous studies [23, 24, 29, 30]. In the study conducted by Galarza *et al*. the sensitivity for detection of RIF resistance was found to be 98.7 % and the specificity 97.75 % [24]. Likewise, based on a meta-analysis in the year 2013, a pooled sensitivity of the high-resolution melting curve analysis was 94%, and the pooled specificity was very high at 99% [23]. One potential reason for a lower sensitivity of HRM in other studies is that 5% of RIF resistance is due to mutations in DNA other than the *rpoB* region [31]. However, in our study, the obtained 100% sensitivity may be due to the fact that none of the RIF-resistant isolates had mutations outside the *rpoB* core.

Our results also indicated that the specificity of the HRM was slightly lower than those found in previous studies [23, 24, 29, 30]. The possible reasons for false-positive results might be as follows: it is possible that these mutations are silent or confer low-level resistance, and it may also occur due to the cross-contamination. In recent years, several rapid molecular assays including commercial kits and Xpert MTB/RIF have been used for the diagnosis of drug resistance in *M*. *tuberculosis* [32–35]. Although rapid, these commercial methods are too expensive and technically demanding in limited-resource countries like Iran.

In the case of HRM, there are advantages over current molecular techniques. For example, the assay is able to determine the RIF resistance in a large number of specimens simultaneously, within approximately 2h of obtaining DNA. Furthermore, the HRM is able to detect mutations without the need of labelled primers, product fractionation, DNA restriction or specific sequence analyses, providing options as to how to meet the needs of the laboratory [17]. An additional appealing feature of HRM is its adaptability. Commercial molecular techniques (i.e. real-time PCR and HRM) cannot detect drug resistance conferred by unknown mutations, and this limitation is common among all of them. However, by HRM, as new mutations are identified, additional probes could easily be designed and added to the assay [17]. This is an important feature because patterns of drug resistance can vary widely in a particular geographic area.

However, results from melting analysis depend on several factors. An important disadvantage of the HRM assay is that the accuracy of the assay critically depends on the sample source, preparation, quality of the extracted DNA, concentration, amplicon length, GC content, equipment, and the dye [22, 23, 36]. The previous study showed that a higher sensitivity could be achieved when fresh samples were used for HRM because the DNA might have degenerated during sampling [22]. Wittwer *et al*. indicated that the method is more sensitive for fragments smaller than 600 bp [37]. Likewise, Taylor *et al*. described in their study that lower GC content might have an association with false-negative results [38]. Moreover, dye and instruments are also very important in the accuracy of HRM. All of these parameters may limit the extensive application of HRM to some degree. Furthermore, standard optimization procedures must be established before HRM is used in clinical laboratories worldwide [23]. Human intervention required with HRM also can increase the risk of contamination.

Our systematic review had some limitations. First, HRM was performed only on culture isolates. If this method could be used for the detection of RIF resistance in *M*. *tuberculosis* from clinical specimens, the diagnostic time would be further shortened. Second, HRM assay may detect silent or other mutations which are not actually associated with drug resistance. Third, we did not study all kinds of mutations related to RIF resistance, and thus, the technique cannot yet replace conventional DST. Fourth, our sample size was low, and at a higher sample size; there would be an increased chance of finding non-*rpoB* mutated resistant strains. In conclusion, the analysis has demonstrated that the diagnostic accuracy of HRM tests is insufficient to replace Xpert MTB/RIF and conventional DST. HRM test may be used in combination with culture due to the advantage of the time to result. Evaluation of new molecular DST would be an important next step to further expedite drug resistant-TB detection.

## Acknowledgment

This study was supported by Shahid Beheshti University of Medical Sciences, Tehran, Iran.

